# Sex Differences in Bed Nucleus of the Stria Terminalis Response to Calcitonin Gene-Related Peptide

**DOI:** 10.64898/2026.05.21.726958

**Authors:** Rebecca Lorsung, Yadong Ji, Nathan Cramer, Rebecca Aitken, Sung Han, Radi Masri, Asaf Keller

**Author notes:** Conflict of Interest Statement: The authors declare they have no conflicts of interest to disclose.

## Abstract

Women are disproportionately affected by chronic pain, yet the neural mechanisms underlying sex differences in affective pain processing remain incompletely understood. The bed nucleus of the stria terminalis (BNST), a sexually dimorphic structure implicated in aversion and chronic pain, receives dense input from aversive calcitonin gene-related peptide (CGRP)–expressing neurons arising from the parabrachial nucleus (PBN). Although CGRP signaling has been implicated in sex differences in clinical pain conditions, whether CGRP transmission within the PBN→BNST pathway is sexually dimorphic has not been determined. Here, we tested the hypothesis that CGRP signaling in the BNST differs between sexes. Contrary to our prediction, PBN CGRP neurotransmitter release in the BNST was sex-independent. However, CGRP neuromodulation of BNST excitability exhibited sex-dependent features. While CGRP potentiated PBN→BNST glutamatergic signaling in both sexes, spontaneous inhibitory signaling was selectively increased in males. Together, these findings indicate that sex differences in this circuit arise not from differential peptide release, but from downstream modulation of inhibitory tone, biasing female BNST neurons toward greater excitation. Such circuit-specific sex differences may contribute to the enhanced susceptibility of females to affective components of chronic pain and highlight targets for sex-informed therapeutic interventions.

## Introduction

Women experience chronic pain more frequently, with greater severity and longer duration than men, across a wide range of conditions (Mogil, 2021, 2012, 2009; Osborne and Davis, 2022). Although environmental and psychosocial factors contribute to these sex differences (Bartley and Fillingim, 2013; Fillingim, 2000), growing evidence highlight biological mechanisms—including genetic, neuroimmune, and neurobiological pathways—as key contributors (Stratton et al., 2024). Identifying the neural circuits that mediate sex-specific differences in chronic pain is essential for developing more effective, sex-informed diagnostics and treatments.

A candidate brain region is the bed nucleus of the stria terminalis (BNST), which is increasingly recognized for its role in affective pain processing and the transition to chronic pain (Deyama et al., 2007; Fang et al., 2023; Minami, 2019; Yu et al., 2021). In animal models, BNST neurons are robustly activated by aversive stimuli (Aston-Jones et al., 1999a, 1999b; Butler et al., 2016; Demaestri et al., 2019; Fang et al., 2023; Goode et al., 2019; Hu et al., 2025; Janitzky et al., 2015). BNST exhibits sex differences in morphology (Aston-Jones et al., 1999a), neuropeptide expression (Uchida et al., 2019; Van Leeuwen et al., 1985), and stress and anxiety-related behaviors (Lebow and Chen, 2016), including sex-dependent responses during aversive processing (Jaramillo et al., 2020; Urien and Bauer, 2022; Yu et al., 2021).

During aversive processing, BNST activity is modulated by the parabrachial nucleus (PBN) (Jaramillo et al., 2020), a region central to the affective dimension of pain (Chiang et al., 2019). A subpopulation of PBN neurons that express calcitonin gene related peptide (CGRP) form large, presumably highly efficient, perisomatic synapses in the dorsolateral/anterolateral BNST (Kozicz and Arimura, 2001; Ye and Veinante, 2019). These PBN_CGRP_ neurons are activated during chronic pain (Chiang et al., 2020), and activation of this pathway is aversive ((Bowen et al., 2020; Jaramillo et al., 2020; Lynch et al., 2024)). Although PBN_CGRP_ neurons co-release glutamate and CGRP (Shimada et al., 1985), a role for CGRP signaling in BNST-mediate aversion is supported by direct infusion studies demonstrating its anxiogenic effect (Kelly S. Sink et al., 2013; K.S. Sink et al., 2013; Sink et al., 2011).

CGRP signaling exhibits pronounced sex differences in pain-related conditions. Clinical and preclinical studies demonstrate enhanced CGRP-dependent pain mechanisms in females (Avona et al., 2019; de Vries Lentsch et al., 2021; Labastida-Ramírez et al., 2019; Paige et al., 2022; Presto and Neugebauer, 2022). Consistent with this, our work and others’ have shown that CGRP exerts sex-dependent effects on neuronal activity in other brain regions (Lorsung et al., 2025; Paige et al., 2022). Despite this, most studies of the PBN_CGRP_→BNST pathway have been conducted exclusively in male animals. A notable exception showed that activation of BNST neurons receiving PBN input increases anxiety-like behavior selectively in females, though CGRP-specific contributions were not examined (Jaramillo et al., 2020). Thus, whether CGRP signaling within the PBN→BNST pathway exhibits sex-dependent effects remains unknown.

Here, we tested the hypothesis that CGRP signaling in the BNST is sexually dimorphic. Because sex differences could arise from differences in peptide release or from differential postsynaptic responsiveness, we assessed both PBN-derived CGRP release and CGRP-induced modulation of BNST neuronal activity. We predicted that females would exhibit greater CGRP release in response to aversive stimuli and that CGRP signaling would produce greater BNST activation in females compared to males. Given that BNST hyperexcitability is associated with chronic pain states (Huang et al., 2021; Lynch et al., 2024; Maruyama et al., 2018; Minami, 2019; Minami and Ide, 2015; Song et al., 2020; Yamauchi et al., 2022), and that sex differences in BNST excitability have been reported in aversive processing (Jaramillo et al., 2020; Song et al., 2020; Yu et al., 2021), such differences in CGRP signaling may contribute to sex-specific vulnerability to chronic pain.

## Methods

### Animals

All procedures adhered to the Guide for the Care and Use of Laboratory Animals and approved by the Institutional Animal Care and Use Committee at the University of Maryland School of Medicine. We used 107 CGRP (calcitonin-gene-related peptide)-CRE heterozygous mice (54 female, 53 male) that were bred in-house from male B6.Cg-Calcatm1.1(cre/EGFP)Rpa/J (stock #033168) x female C57BL/6J mice (strain #000664). Breeding pairs were obtained from The Jackson Laboratory. Offspring were weaned at postnatal day (PD)21 and housed two to five per cage in single-sex groups. Food and water were available ad libitum, and lights were maintained on a 12/12 h light/dark cycle. 8 males and 8 females were used for fiber photometry experiments. The remaining mice (46 female, 45 male) were used for *in vitro* electrophysiology, where 1 to 7 neurons were recorded in each mouse from 1 to 3 BNST slices. In slices treated with CGRP or a CGRP antagonist, only one neuron was recorded per slice. Additionally, no more than 1 to 2 neurons per animal were included in any given experiment to prevent a single animal from disproportionately contributing to the dataset.

### Virus injection

We anesthetized animals with isoflurane and placed them in a stereotaxic frame. Either left or right PBN (-5.2 mm AP, ±1.5 mm ML, -2.9 mm DV) was targeted via a small craniotomy (∼1-2 mm). Only the right PBN was targeted in photometry recordings. We injected 0.5 μL of adeno-associated virus generated by the University of Maryland School of Medicine’s Viral Vector Core – Baltimore, Maryland; AAV5-DIO-ChR2-eYFP, OR 0.25 μL AAVDJ-DIO-CybSEP2. CybSEP2 is a presynaptic pH-sensitive presynaptic sensor which is trafficked by peptide-containing large dense core vesicles (LDCVs), and which undergoes a shift in fluorescence upon LDCV fusion and neuropeptide release (Kim et al., 2024). Viruses were injected using a MICRO2T SMARTouch™ controller and Nanoliter202 injector head (World Precision Instruments) at a flow rate of 100 nL/min. The pipette was left in place for 10 min before being slowly retracted over 5 to 10 min. Mice were given Rimadyl for postoperative analgesia, and allowed 6-10 weeks for virus expression. Injection sites were verified by visually confirming robust eYFP fluorescence in the external PBN.

### *In vitro* slice electrophysiology

Adult mice (2 to 12 months old; median 7 = months) were deeply anesthetized with ketamine/xylazine, and 300 µm-thick coronal brain slices containing the BNST were prepared using a modified slice collection method (Ting et al., 2014). Slicing was performed in ice-cold (4 °C) NMDG-based cutting solution containing: 92 mM N-methyl-D-glucamine (NMDG), 30 mM NaHCO₃, 20 mM HEPES, 25 mM glucose, 5 mM sodium ascorbate, 2 mM thiourea, 1.25 mM NaH₂PO₄, 2.5 mM KCl, 3 mM sodium pyruvate, 10 mM MgSO₄·7H₂O, and 0.5 mM CaCl₂·2H₂O. Slices were incubated in this solution at 37 °C for 7 minutes before being transferred to artificial cerebrospinal fluid (ACSF) at 37 °C for an additional 10–20 minutes, after which they were held at room temperature (25 °C).

For recordings, we placed slices in a submersion chamber continuously perfused (2 mL/min) with ACSF: 119 mM NaCl, 2.5 mM KCl, 1.2 mM NaH_2_PO_4_, 2.4 mM NaHCO_3_, 12.5 mM glucose, 2 mM MgSO_4_·7H_2_O, and 2 mM CaCl_2_·2H_2_O. Both NMDG-based cutting solution and ACSF were adjusted to a pH of 7.4, mOsm of 305, and bubbled with carbogen (95% O2 and 5% CO2) during use and for at least 30 minutes prior to experiments.

We obtained whole-cell voltage-clamp recordings (−70 mV) from the anterolateral BNST using borosilicate pipettes with an impedance of 4 to 6 MΩ. Our internal solution contained the following. For EPSC recordings (low chloride internal solution): 130 mM cesium methanesulfonate, 10 mM HEPES, 1 mM magnesium chloride, 2.5 mM ATP-Mg, 0.5 mM EGTA, 0.2 mM GTP-Tris, 5 mM QX-314, and 2% biocytin (pH of 7.3, 285 mOsm). For IPSC recordings (high chloride internal solution): 70 mM potassium gluconate, 60 mM potassium chloride, 10 mM HEPES, 1 mM magnesium chloride, 2.5 mM Mg-ATP, 0.5 mM EGTA, and 0.2 mM GTP-Tris (pH 7.3, 285 mOsm).

Electrical stimulation for evoked IPSCs was delivered through a bipolar electrode directly dorsal to the recorded cell, to produce a single-component IPSC with amplitude >20 pA. Excitatory postsynaptic currents were optically evoked (oEPSCs) by whole field illumination of the BNST ipsilateral to PBN with prior ChR2 injection at 470 nm (Lambda LS light source, Sutter Instrument) and maximum power of 1.4 mW. Optical stimulation parameters, high frequency stimulation (10 Hz, 3 ms pulse duration) and single/paired exposures (3 ms pulse duration, 100 ms interval) were controlled by a SmartShutter system (Sutter Instrument). We monitored series resistance throughout recordings by measuring the current evoked by a -5 mV square pulse at ∼ 20 s intervals. Evoked oEPSC and IPSC amplitudes were quantified using Clampfit 11.2 (Molecular Devices). Spontaneous EPSCs and IPSCs were quantified using Easy Electrophysiology (Easy Electrophysiology Ltd).

### *In vivo* LDCV quantification

We anesthetized animals using urethane (2 mg/kg) and placed them in a stereotaxic frame. A fiber optic probe (400 μm diameter, 0.39 NA; RWD Life Science) was lowered into the anterolateral BNST (0.14 mm AP, +0.9 mm ML) of adult mice previously injected with AAVDJ-DIO-CybSEP2 (Kim et al., 2024) in the ipsilateral parabrachial nucleus. All CybSEP2 recordings were performed in the right BNST. Optimal depth of recording was determined by observing baseline fluorescence changes and the response to intermittent delivery of 485 μA foot shock (DV: median = 3.835, min = 3.300, max = 4.220). Once the optimal depth of recording was determined, responses to all aversive stimuli (foot shock, tail pinch) were recorded at that depth. Following recordings, mice were deeply anesthetized with ketamine/xylazine, perfused with 4% neutral buffered formalin, and injection site was validated histologically.

For recordings, the fiber optic probe was connected to an RZX10 LUX fiber photometry processor running Synapse software (Tucker-Davis Technologies) through a Doric Mini Cube (Doric Lenses). LEDs at 465 nm and 405 nm were used for CybSEP2 excitation and isosbestic control, respectively. LED power was calibrated using a digital optical power meter (Thorlabs).

We analyzed the data using customized Python scripts adapted from Tucker-Davis Technologies templates which calculated relative changes in fluorescence. These were calculated by subtracting the scaled isosbestic signal (405 nm) from the sensor fluorescence (465 nm). Event related changes in sensor fluorescence were converted to ΔF/F using the 5 second window prior to each stimulation as baseline. The average ΔF/F during the first 10 seconds following aversive stimuli delivery, or from 45-55 seconds, was calculated using Microsoft Excel.

### DAB immunohistochemistry

For generating the example image for Figure 3B, sixty µm thick slices from a female injected with AAV5-DIO-ChR2-eYFP in the PBN, as described above, were washed in 1× PBS (0.145 M NaCl, 0.0027 M KCl, 0.008 M Na₂HPO₄, 0.0015 M KH₂PO₄, pH 7.4) three times for 10 minutes. Tissue sections were incubated in blocking buffer (1% BSA, 0.3% Triton X-100 in 1× PBS) for 1 hour at room temperature to reduce non-specific binding. Primary antibody (1:500 GFP Polyclonal Antibody, Catalog #A10262, Thermo Fisher Scientific, Waltham, MA) diluted in incubation buffer (1% BSA, 0.1% Triton X-100 in 1× PBS) was applied and incubated for 48 hours at 4 °C. Sections were then washed three times for 15 minutes each in wash buffer and then incubated with secondary antibody (1:300 Goat Anti-Chicken IgY Antibody (H+L), Biotinylated, Vector Laboratories, Newark, CA) diluted in the same incubation buffer for 1 hour at room temperature in the dark. Sections were washed again three times for 15 minutes, mounted, and imaged using Mica WideFocal Microhub Automated Microscope (Leica Microsystems, Wetzlar, Germany).

### Experimental Design and Statistical Analysis

Statistical tests were conducted using Prism 10 (GraphPad) or R, and sample size was determined *a priori* using G*Power software suite (Heinrich-Heine, Universität Düsseldorf). Parametric tests were used when appropriate assumptions were met; otherwise, we used nonparametric tests. Averages described in the text are formatted as mean ± SD unless otherwise stated. All figures were designed using a combination of Prism 10 (GraphPad) and Inkscape 1.3.2. Atlas images for Figure 5A were adapted from (Franklin and Paxinos, 2008), and accessed via a web based tool (https://labs.gaidi.ca/mouse-brain-atlas/).

LDCV photometry data in response to tail pinch and foot shock stimulation were pooled for quantification using functional linear mixed modeling as described in Loewinger et al. (2024).

To visualize cumulative inter-event interval and amplitude distributions for spontaneous IPSCs and EPSCs, events were binned using the Freedman–Diaconis rule, in which bin width is calculated as 2 × IQR × n×(-1/3), and a simple heuristic of √n was used to determine the number of bins in the histogram. The interquartile range (IQR) was calculated by averaging the 25th and 75th percentiles across all neurons. The number of events (n) was defined as the maximum number of events recorded for any individual neuron in the dataset, ensuring that bin widths were appropriate for the most active neurons. Bin widths and edges were then applied consistently across all neurons to allow direct comparisons of histogram distributions using Kolmogorov–Smirnov tests in Prism 10 (GraphPad).

## Results

### Aversive shock induces sex-independent parabrachial neuropeptide release in the BNST

While low-frequency firing of PBN_CGRP_ neurons primarily promotes glutamate signaling, high-frequency firing triggers the fusion of large dense-core vesicles (LDCVs), leading to the release of stored neuropeptides (Kim et al., 2024; Lorsung et al., 2025; Qiu et al., 2016; Schöne et al., 2014; Tallent, 2008). PBN_CGRP_ neurons fire at high frequencies in response to aversive input, especially in chronic pain conditions (Raver et al., 2020; Smith et al., 2023; Uddin et al., 2018). However, the kinetics of PBN_CGRP_ neuropeptide release in BNST in response to aversive input are unknown, as neuropeptides are difficult to directly measure *in vivo* (Girven et al., 2022). It is also unknown whether there are sex differences in PBN_CGRP_ neuropeptide release in the BNST. To address this gap in knowledge, we measured PBN_CGRP_ neuropeptide release in the BNST in response to acute aversive stimuli, and compared this response between sexes.

To do this, we expressed the genetically encoded LDCV sensor CybSEP2 in PBN_CGRP_ neurons (Fig. 1A) and performed fiber photometry recordings in the ipsilateral BNST of lightly anesthetized animals. CybSEP2 localizes to LDCVs and undergoes a pH-dependent increase in fluorescence upon vesicle fusion, enabling real-time monitoring of neuropeptide release (Kim et al., 2024). Although 8 mice of each sex were initially recorded, one female was excluded from the final analysis due to the absence of observable LDCV release following aversive stimulation (tail pinch and foot shock), likely attributable to a shorter viral incubation period (2 weeks less than other subjects) at the time of recording. One male was excluded due to markedly atypical release kinetics (Fig. 1C, inset), characterized by a prolonged response (∼3 minutes vs. ∼10 seconds in other mice), with no identifiable technical or anatomical differences.

**Figure 1:**
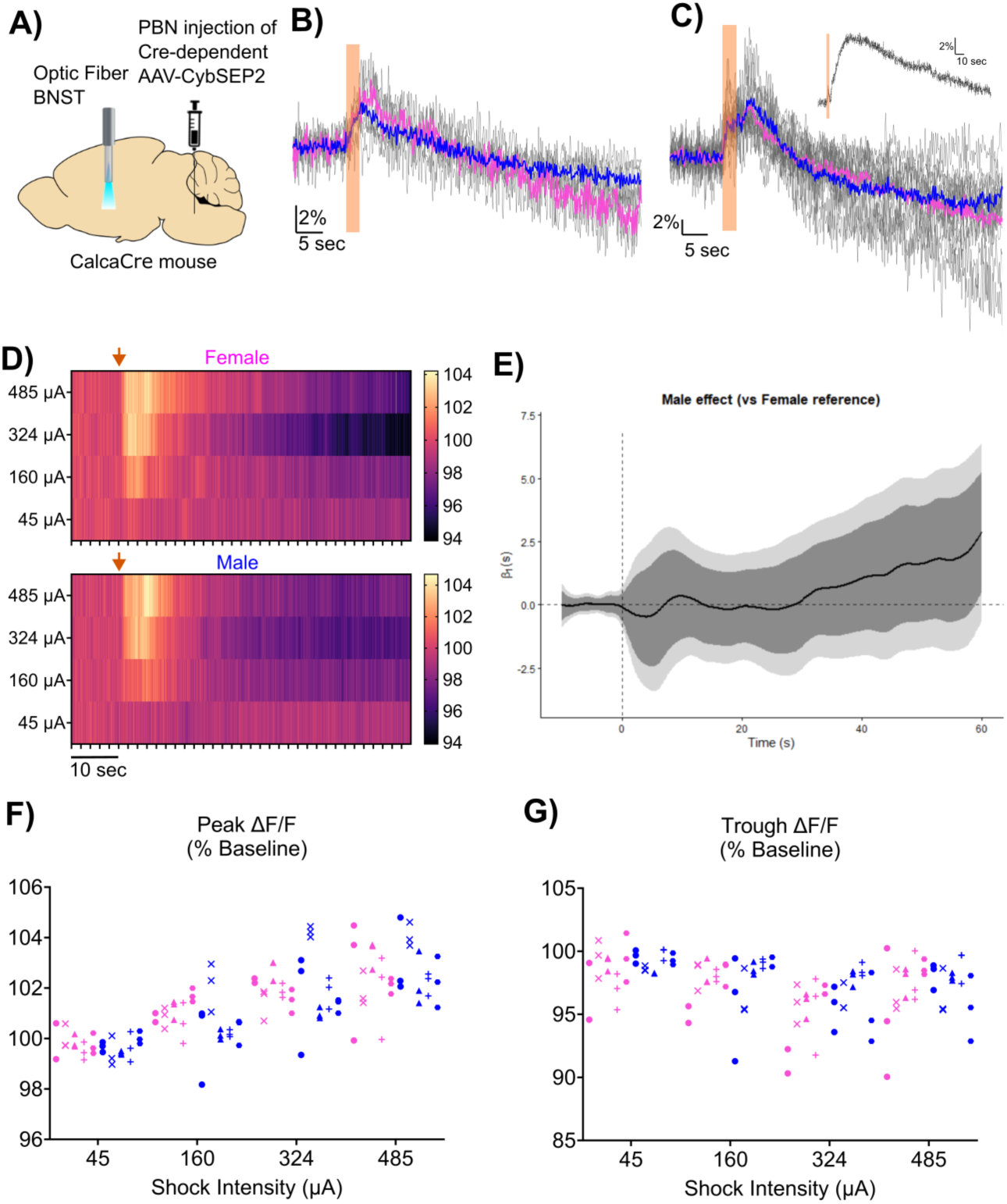
Aversive stimuli induce PBN_CGRP_ LDCV release in BNST. A) Cre-dependent CybSEP2 neuropeptide release sensor was injected into the right lateral parabrachial nucleus (PBN) of male and female Calca_Cre_ mice. Fiber photometry was performed in the ipsilateral BNST of lightly anesthetized mice to monitor peptide release. B–C) ΔF/F traces over time (gray) in response to tail pinch (B, n = 3 f, 4 m) and 485 µA foot shock (D, n = 7 f, 7 m) show transient increases in signal followed by delayed suppression. Multiple trials (2 to 4) were conducted in B and C, and activity was averaged across trials to obtain a single trace per mouse (gray). Average traces are shown for males (blue) and females (pink), while individual animal averages are shown in gray. One male exhibited delayed and prolonged kinetics and is displayed separately as an inset in (C). D) Heat maps showing the average change in ΔF/F over time (% normalized to baseline) in response to different foot shock intensities for pooled trials from all females (top) and males (bottom) that underwent stimulus–response curve recordings (n = 5 females, 5 males). The color bar indicates percentage change in fluorescence from baseline following shock initiation (orange arrow). The black horizontal bar (10 s) indicates the x-axis timescale. E) Estimated functional linear mixed model coefficient over time for sex compared to the female reference. The vertical dashed line marks stimulus onset. Model coefficients are plotted in black, with 95% confidence intervals (CI) shown in gray. Dark gray shading indicates the pointwise CI at each individual time point, while light gray shading indicates the joint CI, which corrects for multiple comparisons across the full time course and is more conservative. F-G) Average normalized ΔF/F magnitude in the first 10 seconds (F) and 45–55 seconds (G) following shock onset remained similar between sexes across shock intensities (n = 5 f, 5 m). Data points represent individual trials from male (blue) and female (pink) mice. Trials from the same mouse are grouped as a column, and share a symbol.

A brief tail pinch (1.5 to 3.6 s) evoked a rapid increase in fluorescence (ΔF/F), consistent with LDCV fusion, followed by a delayed suppression below baseline (Fig. 1B; n = 3 females, 4 males, 2-5 trials averaged per mouse). The response to another aversive stimulus, foot shock (3 s, 485 µA), evoked a similar biphasic response (Fig. 1C; n = 7 females, 7 males, 2 to 3 trials averaged per mouse). In a subset of mice (n = 5 females, 5 males, 2-3 trials averaged per mouse), we generated a stimulus–response curve using a range of shock intensities (45, 180, 360, and 485 µA). The average fluorescence over time for both sexes across all four shock intensities is shown in Figure 1D.

The delayed suppression phase of the fluorescent signal was not attributable to photobleaching. Control recordings without stimulation showed minimal signal drift over comparable time periods. The magnitude of fluorescence decrease during control trials was negligible (−0.002 ± 0.004% ΔF/F), in contrast to the substantially larger suppression observed following shock (−3.34 ± 2.77% ΔF/F). Fluorescence returned to baseline prior to subsequent trials and exhibited a modest increase of 2% across repeated shocks (repeated-measures ANOVA, Treatment effect: F(2, 28) = 6.43, p = 0.02). This slow increase is consistent with activity-dependent potentiation of LDCV release (Kirchner et al., 2023; Park et al., 2006).

To assess whether sex influences LDCV release dynamics, we applied a functional linear mixed model (fLMM) to the full fluorescence time course across all trials in Figure 1B-1D (see Methods, and (Loewinger et al., 2025)). Briefly, the fLMM treats the fluorescence signal as a function over time, and tests whether sex explains variation in the signal beyond baseline variability across animals, enabling the evaluation of potential sex-differences across the time course. In Figure 1E, we visualized the estimated model coefficients over time, where the intercept represents the reference female model and the black line shows the estimated male deviation from this reference across time, with confidence intervals indicated by gray shading. Because the male coefficients overlap with the female intercept throughout the entire time course, these results suggest that the dynamics of LDCV release following foot shock are similar between males and females.

We observed that the strongest responses across the stimulus–response curve in both males and females occurred following 485 µA foot shock (Fig. 1D). This is further illustrated in Figures 1F–G, where average ΔF/F is quantified at two time windows along the biphasic response: during the period of increased fluorescence (Fig. 1F, 1–10 s post-stimulus) and during delayed suppression (Fig. 1G, 45–55 s post-stimulus).

Linear mixed-effects ANOVAs (Type III, Satterthwaite’s method) confirmed a main effect of shock strength but no effect of sex or interaction for either the peak response (Fig. 1F; shock strength F(3, 101.13) = 64.03, p < 2 × 10⁻¹⁶; sex F(1, 8.09) = 0.02, p = 0.88; interaction F(3, 101.13) = 1.34, p = 0.26) or delayed post-shock suppression (Fig. 1G; shock strength F(3, 99.69) = 13.18, p = 2.58 × 10⁻⁷; sex F(1, 7.64) = 0.80, p = 0.40; interaction F(3, 99.69) = 1.03, p = 0.38). Together, these results indicate that LDCV release exhibits a dose-dependent response, in which stronger shocks evoke larger initial increases—consistent with greater LDCV release—and greater delayed suppression in fluorescence. No sex differences were detected across the stimulus–response curve, consistent with the fLMM analysis of the pooled aversive stimuli dataset (Fig. 1E).

We further verified that the reduced responses at lower stimulus intensities were not due to accommodation, as repeated 485 µA shocks delivered at the end of the dose–response series elicited responses comparable to those delivered at the beginning (paired t-test: t = 1.25, p = 0.25; n = 9 mice).

Together, these data demonstrate that aversive mechanical and electrical stimuli reliably evoke PBN_CGRP_ neuropeptide release in the BNST, characterized by a biphasic response consisting of rapid LDCV fusion followed by delayed suppression. The magnitude of both phases scales with stimulus intensity, indicating intensity-dependent recruitment of neuropeptide release mechanisms. We found no evidence for sex differences in release kinetics or magnitude. Overall, PBN_CGRP_ neuropeptide release in response to aversive stimuli appears largely sex-independent.

### CGRP potentiates spontaneous inhibitory signaling only in males

The results described above reject the hypothesis that aversive stimuli evoke different neuropeptide release from PBNCGRP terminals in the BNST of males compared to females. We tested the alternative hypothesis that PBN_CGRP_ neuropeptide release has sex-specific effects on BNST activity. While LDCVs in PBN_CGRP_ neurons may contain several neuropeptides (Missig et al., 2014; Pauli et al., 2022; Shimada et al., 1985; Yamano et al., 1988), we chose to focus on CGRP because of its documented sex-dependent effects (Avona et al., 2019; Lorsung et al., 2025; Paige et al., 2022; Presto and Neugebauer, 2022). CGRP is prominently expressed in PBN_CGRP_ terminals within the BNST, as shown in the super-resolution STED (Stimulated Emission Depletion) image in Figure 2A (reproduced with permission from (Aitken et al., 2026)), which reveals dense CGRP labeling in genetically labeled eYFP+ PBN_CGRP_ terminals, consistent with previous work showing that PBN is the main source of CGRP in the BNST (Dobolyi et al., 2005; Missig et al., 2014; Shimada et al., 1985). We tested whether CGRP exerts a sex dependent effect on inhibitory signaling in the BNST because the majority of synaptic connections between BNST neurons are inhibitory (Sun and Cassell, 1993).

**Figure 2:**
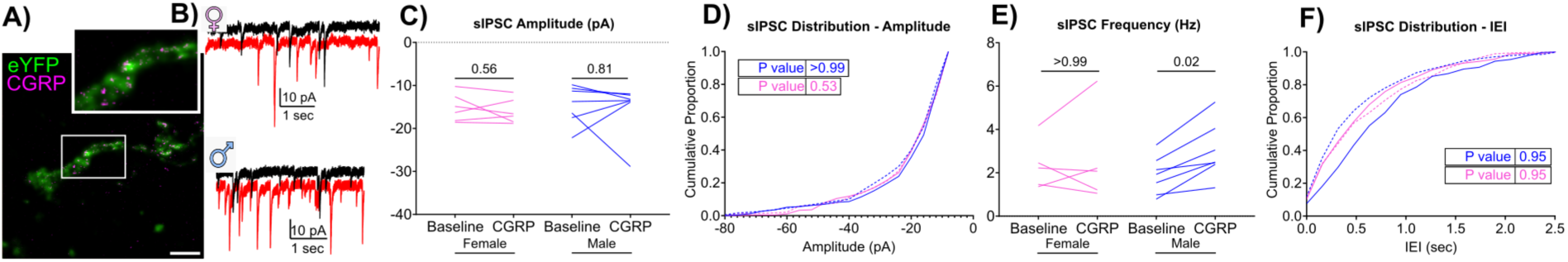
CGRP potentiates spontaneous inhibitory signaling only in males. A) PBN_CGRP_ terminals labeled with a Cre-dependent eYFP virus (green) co-express CGRP (purple), visualized using STED microscopy in the BNST. Inset indicated by white rectangle. Scale bar = 2 µm. B) Example traces of spontaneous inhibitory postsynaptic currents (sIPSCs) recorded from BNST neurons in female (♀) and male (♂) mice. Baseline (black) and 6– to 12 min post–CGRP exposure (500 nM; red) are shown. C) Median sIPSC amplitude was unchanged in both sexes following CGRP exposure. D) Cumulative distributions of sIPSC amplitudes were unchanged from baseline (solid lines) following CGRP application (dashed lines) in both sexes. E) Median sIPSC frequency increased only in male neurons following CGRP exposure. F) Cumulative distributions of sIPSC IEIs were unchanged from baseline (solid lines) following CGRP application (dashed lines) in both sexes. In panels C-F, males are represented in blue and females in pink.

Spontaneous inhibitory postsynaptic currents (sIPSCs) were isolated by recording with a high-chloride internal solution in the continuous presence of CNQX (20 µM) and AP5 (50 µM) to block glutamatergic transmission. To determine the specific effects of CGRP on BNST inhibitory signaling, we bath-applied CGRP (500 nM) to slices, and quantified its effects on sIPSC amplitude and frequency at 6 to 12 minutes of application—an interval previously shown to be sufficient for inducing CGRP-dependent changes in both the rat BNST and the mouse central amygdala (Gungor and Pare, 2014a; Han et al., 2010; Okutsu et al., 2017) compared to a 3 minute baseline recording. Notably, prior studies investigating CGRP signaling in the BNST have focused exclusively on male subjects, leaving open the question of whether CGRP modulates BNST inhibitory transmission in a sex-dependent manner.

Example traces of sIPSCs from a male and a female BNST neuron are shown (Fig. 2B). CGRP application did not affect median sIPSC amplitude in either sex (Fig. 2C; males: p = 0.81, W = –4.00, n = 7 neurons; females: p = 0.56, W = –7.00, n = 6 neurons; Wilcoxon signed-rank test), nor the distribution of sIPSC amplitudes (Fig. 2D; males: p = 0.86, D = 0.15; females: p = 0.53, D = 0.26; Kolmogorov-Smirnov test, binning parameters as described in Methods). These results suggest that CGRP does not alter postsynaptic responsiveness to spontaneous inhibitory inputs in either sex.

However, while female neurons showed no change in median sIPSC frequency (Fig. 2E; p = 0.56, W = –7.00, n = 6 neurons; Wilcoxon signed-rank test), CGRP produced a 29% increase in median sIPSC frequency from 1.93 Hz to 2.49 Hz (Fig. 2E; p = 0.02, W = 28.00, n = 7 neurons; Wilcoxon signed-rank test). Despite this, the cumulative distribution of sIPSC interevent intervals was unchanged in both sexes (Fig. 2F; males: p = 0.95, D = 0.18; females: p = 0.95, D = 0.18; Kolmogorov-Smirnov test, binning parameters as described in Methods). This suggests that CGRP produces a modest increase in sIPSC event frequency without substantially altering the overall distribution of interevent intervals. Limited sampling duration (5 s recordings acquired every 20 s) and low baseline event rates in some neurons (<2 Hz) may have reduced sensitivity to detect subtle distributional differences.

Taken together, these results suggest that CGRP signaling selectively enhances inhibitory activity in males by increasing the frequency of sIPSCs, without impacting the postsynaptic response to inhibitory inputs.

### CGRP has a heterogenous effect on evoked inhibitory tone

As spontaneous IPSCs only capture active inhibitory inputs, they may not capture CGRP’s effects on silent or less-active synapses. To assess whether PBN_CGRP_ neuropeptide release, including CGRP, modulates the broader pool of inhibitory inputs in a sex-dependent manner, we examined the effect of endogenously released PBN_CGRP_ neuropeptides on evoked IPSCs.

To evoke endogenous neuropeptide release in the BNST, we injected a Cre-dependent, channel rhodopsin (ChR2) eYFP-tagged viral construct in the external lateral PBN of CGRP-Cre mice and performed whole cell slice electrophysiology in the BNST (Fig. 3A). Figure 3B depicts a section through BNST from a mouse injected with AAV5-DIO-ChR2-eYFP in the external lateral PBN. Note the dense, perisomatic innervation of labeled, large PBN_CGRP_ presumptive terminals, located primarily in the anterolateral BNST, consistent with previous anatomical reports (Campos et al., 2017; Carter et al., 2013; Chen et al., 2018; Fetterly et al., 2019; Flavin et al., 2014). Guided by the distribution of eYFP-labeled PBN_CGRP_ terminals, we targeted recordings to the ipsilateral anterolateral BNST, where PBN_CGRP_ input was densest. A subset of BNST neurons within this region received direct excitatory input from PBN_CGRP_ terminals, as confirmed by optically-evoked excitatory postsynaptic currents (oEPSCs) in response to a single 470 nm light pulse (3 ms duration, 0.05 Hz). Figure 3C shows a representative oEPSC (black trace), which was suppressed by the AMPA receptor antagonist CNQX (20 μM) and NMDA receptor antagonist AP5 (50 µM) (gray trace). This effect was observed in 32 of 32 neurons, confirming that PBN_CGRP_ inputs to BNST are glutamatergic.

**Figure 3:**
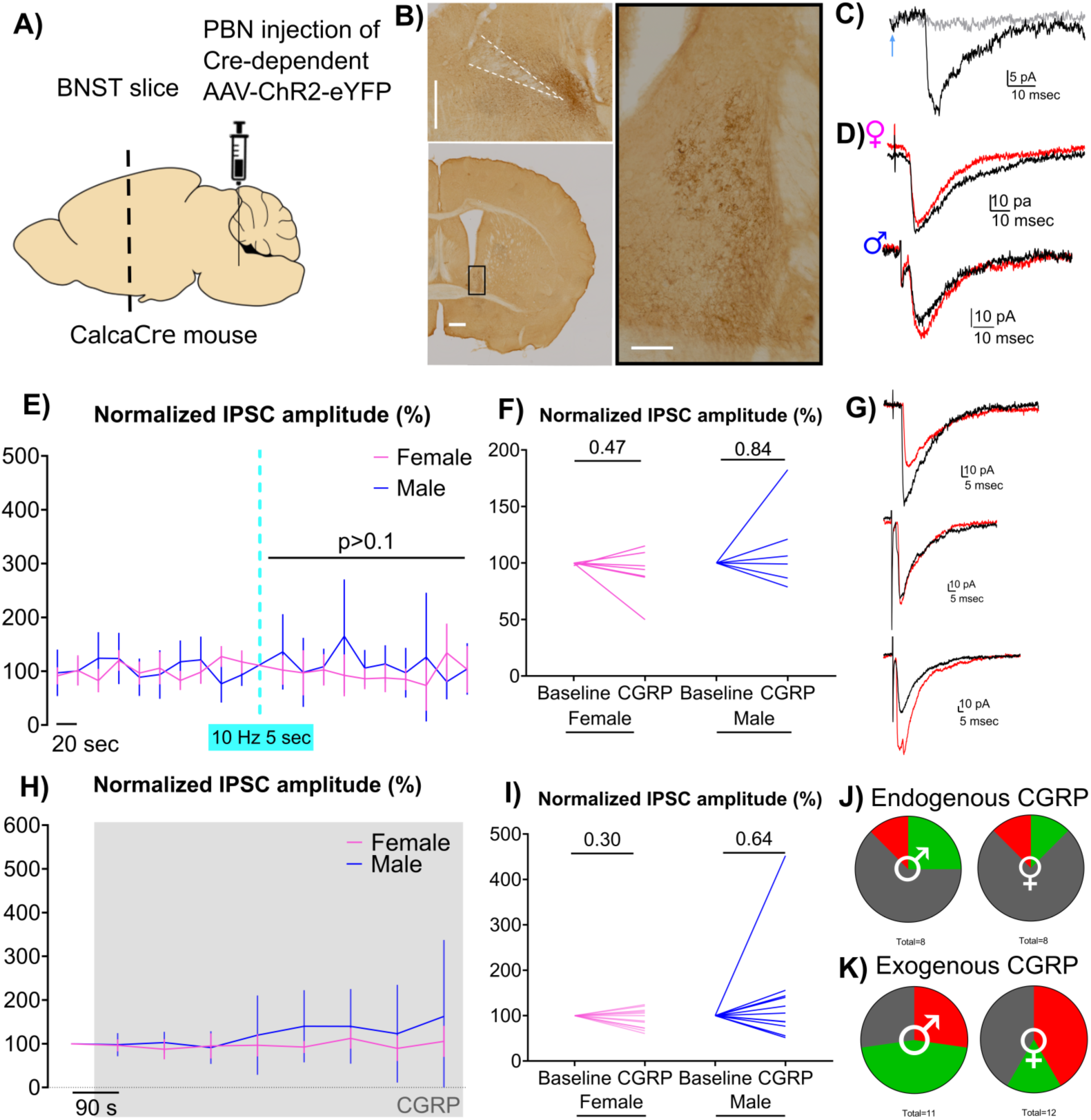
Endogenous CGRP has minimal effect on inhibitory tone, while exogenous CGRP has a heterogeneous effect. A) A Cre-dependent ChR2 virus was injected into the lateral parabrachial nucleus (PBN) of male and female Calca^Cre^ mice. Slice electrophysiology was performed in the ipsilateral BNST. B) Injection site in the PBN (i) and terminal innervation in the BNST (ii) were visualized via DAB immunohistochemistry). (iii) Inset shows dense, perisomatic innervation by large PBN_CGRP_ terminals. Scale bars: i–ii = 500 µm; iii = 100 µm. C) A single 3 msec 470 nm light pulse (arrowhead) evoked an excitatory postsynaptic current (black), which was suppressed by AMPA receptor antagonist CNQX (20 µM; gray). Traces from a representative neuron are shown. Similar effects were observed in 32 of 32 neurons measured. D) Example electrically evoked inhibitory postsynaptic currents (IPSCs) recorded from female (♀, top) and male (♂, bottom) BNST neurons. Recordings were performed in the presence of 20 µM CNQX and 50 µM AP5 to block glutamatergic transmission, using a high-chloride internal solution. Traces show baseline IPSCs (black) and responses following high-frequency optogenetic stimulation (10 Hz, 5 sec; red). All neurons were first confirmed to receive PBN_CGRP_ glutamatergic input prior to CNQX/AP5 application. E) Time course of evoked IPSC amplitude, normalized to baseline, showing no effect of high-frequency stimulation (cyan dashed line). Error bars represent 95% CI. F) No change in normalized IPSC amplitude was observed during the 100 seconds following stimulation in either sex. Pink lines indicate female neurons; blue indicate male neurons. Uncorrected IPSC amplitudes in pA: male; Baseline: −61.76 ± 35.79, CGRP: −68.61 ± 44.90, female; Baseline: −237.1 ± 421.1, CGRP: −220.4 ± 398.8. G) Representative traces of electrically evoked IPSCs recorded from BNST neurons that showed suppression (top), no change (middle) or potentiation (bottom) in IPSC amplitude following application of 500 nM CGRP (red) compared to baseline (black). Recordings were performed in the continued presence of CNQX/AP5 and high-chloride internal. H) Time course of normalized evoked IPSC amplitude showing no consistent effect of 500 nM CGRP application (gray box). Error bars represent 95% CI. I) No change in normalized IPSC amplitude was observed 6 to 12 minutes after CGRP application in either sex. Uncorrected IPSC amplitudes in pA: female; baseline: -69.53 ± 46.02; CGRP: -61.03 ± 35.80, n = 12, male; baseline: -49.94 ± 27.78; CGRP: -65.24 ± 56.68, n = 11. J-K) Proportion of male (♂) and female (♀) neurons showing no change (gray), an increase (green), or a decrease (red) in evoked IPSC amplitude (±20% from baseline) in response to endogenous PBN_CGRP_ neuropeptide release (J, p = >0.99; Fisher’s exact test) or exogenous 500 nM CGRP application (K, p = 0.37; Fisher’s exact test).

We selected BNST neurons receiving direct PBN_CGRP_ excitatory input to test the effect of endogenous neuropeptide release on inhibitory transmission, maximizing the likelihood that released neuropeptides would act locally at the recorded cell. After identifying such neurons, we recorded inhibitory postsynaptic currents (IPSCs) evoked by single-pulse electrical stimulation (1 ms duration, 0.05 Hz) using a stimulating electrode placed dorsal to the recording site within the BNST. To isolate inhibitory responses, we included CNQX (20 µM) and AP5 (50 µM) to block AMPA and NMDA receptors, respectively.

To drive neuropeptide release from PBN_CGRP_ terminals, we applied an optical high-frequency stimulation protocol (10 Hz, 5 sec). While low-frequency firing of PBN_CGRP_ neurons primarily elicits glutamatergic transmission, high-frequency stimulation is necessary to induce endogenous neuropeptide release (Kim et al., 2024; Lorsung et al., 2025; Qiu et al., 2016; Schöne et al., 2014; Tallent, 2008). The amplitude of evoked IPSCs was quantified before and after high frequency optical stimulation to assess the modulatory effects of PBN_CGRP_-derived neuropeptides on inhibitory synaptic input to BNST neurons.

As shown in Figure 3D, endogenously evoked neuropeptide release did not alter electrically evoked IPSC amplitudes in a male or female BNST neuron. This was consistent across both sexes, with no change in amplitude in either sex (Fig. 3E; time: p = 0.73, F(3.449, 26.14) = 0.47; sex: p = 0.19, F(1, 8) = 2.07; interaction: p = 0.27, F(3.449, 26.14) = 1.39; n = 5 male, 5 female neurons; mixed-effects model). Additionally, there was no change in the average post-release IPSC amplitude following stimulation in either sex (Fig. 3F, males: *p* = 0.44, *W* = 7.00, *n* = 5 neurons; females: *p* = 0.31, *W* = –9.00, *n* = 5 neurons; Wilcoxon signed-rank test).

We characterized individual neuronal responses (increase, decrease, no change) to capture heterogeneity that might be obscured by population averages. Due to the limited number of baseline measurements, formal statistical comparisons at the single-cell level were not feasible. Therefore, we applied a descriptive threshold of a 20% change from baseline to classify response direction, providing a qualitative overview of response patterns across the population.

Most BNST neurons (3 of 5 male neurons, 4 of 5 female) maintained IPSC amplitudes within 20% of baseline, while the remaining neurons showed either an increase or decrease in amplitude following high-frequency stimulation (2 of 5 male, 1 of 5 female; Fig. 3J). There was no sex difference in the proportion of BNST neurons that exhibited each response profile following neuropeptide release (p = 0.44; Fisher’s exact test).

There was also no change in access resistance (males: *p* = 0.44, *W* = –9.00, *n* = 6 neurons; females: *p* = 0.47, *W* = –10.00, *n* = 7 neurons; Wilcoxon signed-rank test) in either sex, suggesting that the recording quality remained stable, increasing our confidence that a biological effect was not masked due to a technical change in cell access. Additionally, the holding current remained consistent (males: *p* = 0.16, *W* = –15.00, *n* = 6 neurons; females: *p* = 0.38, *W* = –12.00, *n* = 7 neurons; Wilcoxon signed-rank test) in both sexes; a change in holding current could indicate a subtle change in overall cell excitability or membrane conductance beyond what can be captured via measured IPSC amplitude. Taken together, this suggests that endogenous neuropeptide release has no effect on IPSC amplitude in BNST neurons.

It is possible that endogenously released neuropeptides, induced by high-frequency optic tetanus, are too transient and spatially restricted to alter signaling at inhibitory synapses, which may be distant from the sites of neuropeptide release. To address this possibility, and to specifically examine the effect of CGRP among the neuropeptides co-released by PBN_CGRP_ neurons (Pauli et al., 2022; Sanz et al., 2009), we bath-applied CGRP (500 nM) to investigate the effect of continuous CGRP signaling. This concentration was selected based on prior studies demonstrating inhibitory postsynaptic potential changes in male rats (Gungor and Pare, 2014a), and in mouse spinal cord (Paige et al., 2022).

As seen in Figures 8G-I, the evoked IPSC response to exogenous CGRP was heterogeneous. Sample traces are shown from neurons that potentiated, suppressed, or showed no change (Fig. 3G) in IPSC amplitude following CGRP application. On average, there was no change in IPSC amplitude over time (Fig. 3H; time: p = 0.33, F(1.57, 27.14) = 1.12; sex: p = 0.25, F(1, 21) = 1.41; interaction: p = 0.39, F(1.57, 27.14) = 0.91; n = 11 male, 12 female neurons; mixed-effects model with repeated measures). Similarly, the average IPSC amplitude 6 to 12 minutes after CGRP application, a timepoint sufficient to induce exogenous CGRP-dependent changes in rat BNST (Gungor and Pare, 2014) and mouse central amygdala (Han et al., 2010; Okutsu et al., 2017), showed no change compared to baseline in either sex (Fig. 3I, male: p = 0.64, W = 12; n = 11; Wilcoxon signed-rank test; female: p = 0.30, t = 1.08; n = 12 neurons; paired t-test). There was no change in access resistance (males: p = 0.46, W = -18.00; n = 11 neurons; Wilcoxon signed-rank test; females: p = 0.14; t=1.596, n = 12 neurons; paired t-test), or holding current (males: p = 0.64, W = -12.00; n = 11 neurons; females: p = 0.42, W = -22.00; n = 12 neurons; Wilcoxon signed-rank test) in CGRP-treated neurons.

We characterized the heterogeneity of the population response as described above. A subset of BNST neurons exhibited a >20% increase in IPSC amplitude following 6–12 minutes of CGRP application (5 of 11 male neurons, 2 of 12 female neurons), while other BNST neurons showed a decrease in IPSC amplitude (3 of 11 male neurons, 5 of 12 female neurons) following 6-12 minutes of CGRP application (Fig. 3K). There was no sex difference in the proportion of BNST neurons to exhibit each response profile following CGRP exposure (p = 0.37; Fisher’s exact test). Whether or not CGRP may elicit a change in inhibitory current amplitudes in select BNST neuronal subpopulations (discussed further below) there is no population effect of CGRP on evoked inhibitory signaling in the BNST.

To test whether CGRP alters GABA presynaptic release probability, we examined paired-pulse ratio (PPR) of these evoked IPSCs in a subset of the neurons (Fig. 3H-I). PPR is defined as the ratio of the amplitude of the second to the first response to closely spaced stimuli (150 ms inter-stimulus interval). A change in the PPR would suggest CGRP modulates presynaptic release (Debanne et al., 1996; Dobrunz and Stevens, 1997; Kim and Alger, 2001; Manabe et al., 1993). We compared the average IPSC PPR 6 to 12 minutes after CGRP application to baseline PPR, and found no effect of treatment, sex, or their interaction (treatment: p = 0.63, F(1,10) = 0.25; sex: p = 0.19, F(1,10) = 1.93; interaction: p = 0.83, F(1,10) = 0.05; n = 7 male neurons, 5 female neurons; two-way RM ANOVA). Evoked IPSC PPR values were similar between groups in both at baseline (male: 1.11 ± 0.29, n = 7; female: 0.95 ± 0.22, n = 5) and following CGRP application (male: 1.13 ± 0.16, n = 7; female: 1.01 ± 0.19, n = 5).

These PPR recordings were obtained from the same neurons in which sIPSCs were measured (Fig. 2). Because increases in sIPSC frequency can reflect enhanced presynaptic GABA release probability, we tested whether neurons exhibiting increased sIPSC frequency also showed corresponding changes in PPR by assessing the correlation between CGRP-induced changes in sIPSC frequency and PPR (normalized to baseline). There was no correlation across the pooled set of neurons (r = -0.17, p = 0.60, R² = 0.03, n = 7 male, 5 female neurons, Pearson correlation), nor when restricted to male neurons (r = 0.14, p = 0.76, R² = 0.02, n = 7, Pearson correlation), which exhibited CGRP-induced increases in sIPSC frequency (Fig. 2D). These findings suggest that, even in neurons with increased spontaneous inhibitory event frequency, CGRP does not alter PPR at the electrically evoked synapses tested.

Together, these findings suggest that CGRP-induced changes in spontaneous inhibitory event frequency are not associated with changes in evoked presynaptic GABA release probability. While this does not rule out modulation of presynaptic release probability in inhibitory inputs which were not recruited by the stimulation electrode, an alternative explanation is that CGRP increases the excitability of a local inhibitory neuronal population, as has been observed in the central amygdala (Han et al., 2010).

### Sex-Independent CGRP potentiation of PBN_CGRP_→BNST glutamatergic signaling

CGRP can also modulate glutamatergic signaling. In the central amygdala, CGRP potentiates glutamatergic inputs by acting on NMDA receptors (Han et al., 2010; Okutsu et al., 2017), with our work showing that preferential potentiation in female compared to male neurons (Lorsung et al, 2025). Whether CGRP potentiates glutamatergic signaling in the BNST, and whether there are sex differences in the degree of this potentiation, is not known. Therefore, we compared, in both sexes, the effect of endogenous CGRP release on PBN_CGRP_ glutamatergic inputs to BNST neurons.

To establish a baseline glutamatergic response, optically evoked EPSCs (oEPSCs) were recorded at 0.05 Hz in BNST neurons from CGRP(Calca)_Cre_ mice expressing ChR2 in PBN_CGRP_ terminals. Recordings were obtained using either low-chloride internal solution, which favors detection of inward EPSCs while minimizing inhibitory currents, or high-chloride internal solution. Data from both conditions were pooled for analysis, as baseline oEPSC amplitudes did not differ between internal solutions (two-tailed Mann–Whitney U test, U = 880.5, exact p = 0.74; Low chloride internal solution: median = −17.13 pA, n = 46; high chloride internal solution: median = −13.41 pA, n = 40).

This baseline oEPSC measurement was followed by a high-frequency optical stimulation protocol (10 Hz for 5 seconds) to induce neuropeptide release. After stimulation, single-pulse oEPSCs were recorded again to assess changes in glutamatergic transmission following peptide release.

Figure 4A depicts oEPSCs recorded from male and female BNST neurons before and after neuropeptide release, showing an 82% and 99% increase in the amplitude of the glutamatergic response following tetanus stimulation, respectively. Quantification of the kinetics of the group response (Fig. 4B) revealed an effect of time, but no effect of sex or interaction (time: p < 0.0001, F(6.23, 408.8) = 7.41; sex: p = 0.98, F(1, 69) = 0.0004; interaction: p = 0.14, F(6.23, 408.8) = 1.62; N=42 male, 34 female neurons; mixed-effects model with repeated measures). Post hoc comparisons revealed that oEPSCs remain potentiated during the first 100 seconds following high-frequency stimulation compared to baseline (Uncorrected Fisher’s LSD). The average post-tetanus response is quantified over this 100-second period in Figure 4C. While there is an effect of condition (Baseline vs. CGRP), there was no effect of sex or interaction between sex and condition (condition: p = 2.57e-05, F(1, 80.1) = 19.96; sex: p = 0.32, F(1, 73.0) = 1.02; interaction: p = 0.31, F(1, 80.1) = 1.05; mixed-effects model). Post-hoc analysis confirmed that CGRP increased oEPSC amplitude relative to baseline in both females and males (Fig. 4C; females: p = 0.0022, t = -3.69, mean diff. = 38.4%; males: p = 0.05, t = -2.57, mean diff. = 24.1%; Tukey-adjusted t-test), but there was no difference in oEPSC amplitude between males and females at the CGRP condition (p = 0.49, t = 1.43; Tukey-adjusted t-test). Additionally, the magnitude of oEPSC potentiation did not differ between internal solutions (two-tailed Mann–Whitney U test, U = 794.5, exact p = 0.58; low-chloride: median = 1.159, n = 45; high-chloride: median = 1.228, n = 38).

**Figure 4:**
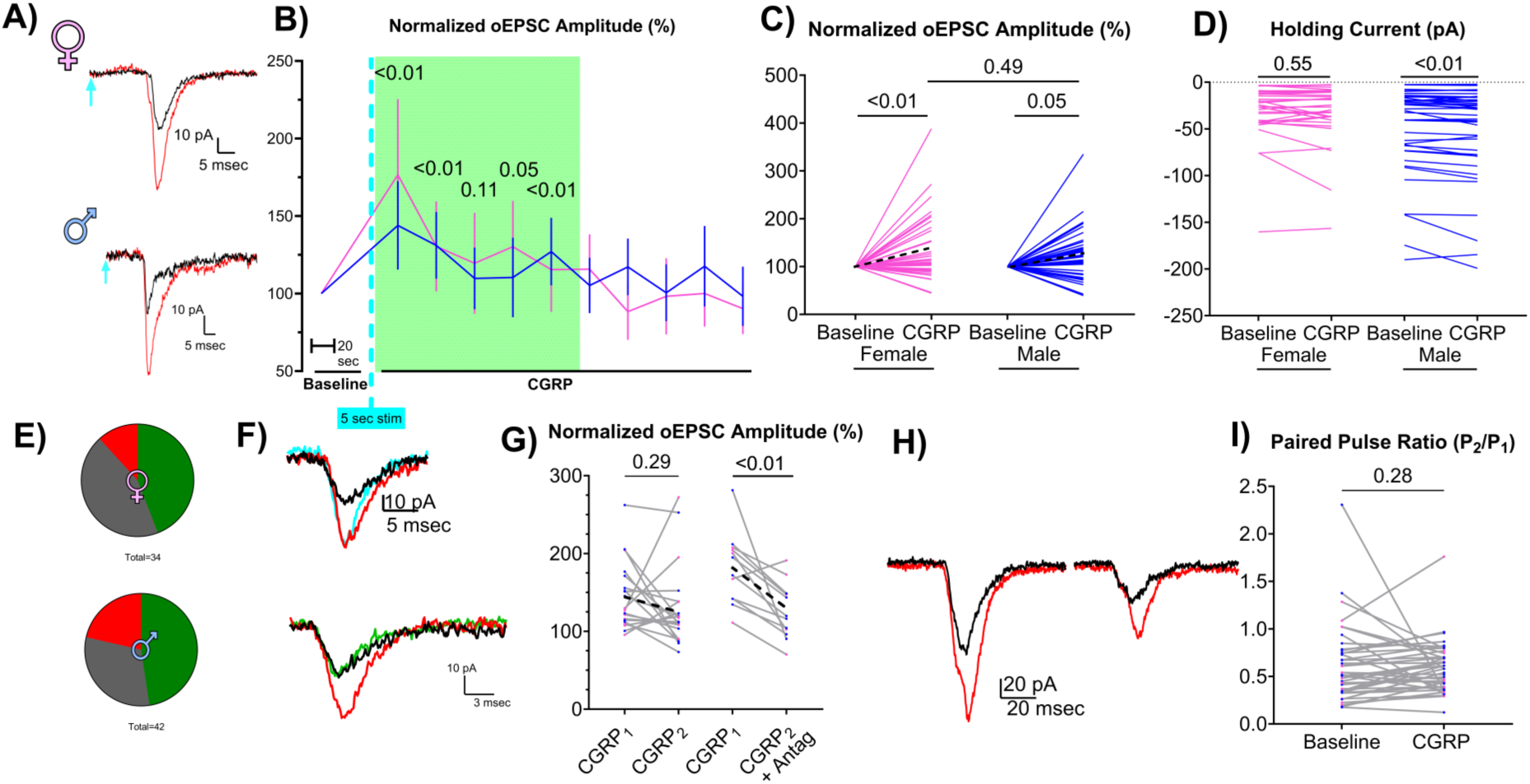
Endogenously released CGRP evokes transient, postsynaptic glutamatergic potentiation in both male and female BNST neurons. A) Representative optically evoked excitatory postsynaptic currents (oEPSCs) recorded from male (♂) and female (♀) BNST neurons before (black) and after high-frequency optical stimulation (10 Hz, 5 s, red) show potentiation. B) Time course of normalized oEPSC amplitude following stimulation (cyan dashed line), plotted by sex. Green box highlights the time window (0 to 100 s post-stimulation) during which oEPSC potentiation was observed in both sexes. Error bars represent 95% confidence intervals. C) Average oEPSC amplitude during the 0 to 100 s window post-stimulation, showing similar levels of potentiation in males and females. D) Baseline holding current decreases only in male BNST neurons during this post-stimulation period. E) Proportion of BNST neurons showing an increase (green), decrease (red), or no change (gray) in oEPSC amplitude (±20% from baseline) was similar between sexes. (p = 0.38, χ²(2) = 1.96; chi-square test) F) Top - Representative oEPSCs from a BNST neuron showing similar levels of potentiation following two consecutive high-frequency optical stimuli (10 Hz, 5 s; red and cyan traces) compared to baseline (black). oEPSC amplitude returned to baseline levels prior to delivery of second stimulus. Bottom - In another neuron, the first optical stimulus induces oEPSC potentiation (red), but a second stimulus delivered in the presence of 1 µM CGRP receptor antagonist (CGRP₈₋₃₇) fails to induce potentiation (green). G) Group data showing no difference in potentiation amplitude between the first and second tetanus delivered (CGRP1 vs. CGRP2), but a reduction in potentiation when the second stimulus is delivered in the presence of CGRP antagonist (CGRP2 + Antag). H) Example oEPSCs evoked by paired-pulse stimulation (inter-stimulus interval: 150 msec), showing comparable potentiation of both pulses following high-frequency optical stimulation (red) relative to baseline (black). I) Group data showing no change in paired-pulse ratio (PPR: pulse 2 / pulse 1 amplitude) in the 0–100 s period following stimulation compared to baseline. Pink and blue lines or datapoints represent female or male neurons, respectively.

There was a slight accompanying drop in the holding current in males in the 100 seconds following high frequency stimulation (Fig. 4D, p < 0.01, W = –488.0, median diff. = –1.93 pA; n = 39 neurons; Wilcoxon signed-rank test), but not in females (Fig. 4D, p = 0.55, W = –66.0; n = 32 female neurons; Wilcoxon signed-rank test). There was no change in access resistance in either males (p = 0.33, N = 46 neurons from 28 male mice, Wilcoxon test) or females (p = 0.64, N = 35 neurons from 26 female mice, Wilcoxon test), suggesting similar voltage control across sexes before and after PBN_CGRP_ neuropeptide release. These data suggest that neuropeptide release from PBN_CGRP_ inputs transiently potentiates PBN_CGRP_ glutamatergic signaling in the BNST in both sexes, while there is a small accompanying change in membrane ion conductance only in males.

Although there was variability in the post-tetanus response amplitude and direction across neurons (Fig. 4E), nearly half of the neurons in both sexes exhibited potentiated oEPSC amplitudes greater than 120% of baseline (20 of 42 male neurons, 15 of 34 female neurons). A minority showed oEPSC suppression below 80% of baseline (9 of 42 male neurons, 4 of 34 female neurons), while some neurons displayed no change, remaining within 20% of baseline (13 of 42 male neurons, 15 of 34 female neurons). There were no sex differences in the distribution of these response profiles following high-frequency stimulation (Fig. 4E, p = 0.38, χ²(2) = 1.96; chi-square test). This suggests that there is a heterogeneity of response profiles in the BNST following PBN_CGRP_ neuropeptide release across both males and females. Given that there were no sex differences in response amplitude (Fig. 4C), kinetics (Fig. 4B), or response types (Fig. 4E), male and female neurons were grouped together for subsequent analysis examining mechanism.

While high-frequency optic tetanus induces LDCV release (Kim et al., 2024; Lorsung et al., 2025; Qiu et al., 2016; Schöne et al., 2014; Tallent, 2008), PBN_CGRP_ LDCVs co-express neuropeptides other than CGRP (Pauli et al., 2022; Sanz et al., 2009). We tested whether potentiation of PBN_CGRP_ glutamatergic signaling following high-frequency stimulation is caused specifically by CGRP release, as has been previously observed in the central amygdala (Han et al., 2010; Lorsung et al., 2025; Okutsu et al., 2017). As shown in Fig. 4F, a second series of PBN_CGRP_ terminal high-frequency stimulation produces a similar potentiation magnitude as an initial delivery. However, application of the CGRP receptor antagonist CGRP_8-37_ (1 μM) prevented potentiation. This is quantified in Figure 4G, where a second stimulation produces a similar degree of potentiation (p = 0.29, t = 1.08, mean diff. = –14.09%; n = 20 neurons; paired t-test), while CGRP_8-37_ suppresses this second potentiation (p < 0.01, W = –76.0, median diff. = –48.60%; n = 12 neurons; Wilcoxon test). These findings suggest that CGRP signaling plays a role in enhancing PBN_CGRP_→BNST glutamatergic signaling following PBN_CGRP_ neuropeptide release.

We investigated the mechanism of CGRP signaling by testing whether CGRP-dependent glutamate potentiation is mediated through a pre- or postsynaptic mechanism. To assess this, we measured paired-pulse oEPSC amplitudes before and after high-frequency optic tetanus. An example neuron shown in Figure 4H demonstrates similar potentiation in both the first and second oEPSCs, resulting in no change in PPR following high-frequency stimulation. As quantified in Figure 4I, there was no change in oEPSC PPR across recorded neurons (p = 0.28, W = 169.0; n = 41 neurons; Wilcoxon test). This suggests that CGRP is acting through a postsynaptic mechanism in the BNST.

Taken together, these data suggest that while there is heterogeneity in individual neuronal responses, both males and females exhibit similar transient postsynaptic potentiation of PBN_CGRP_ glutamatergic signaling in response to CGRP release in the BNST.

We next asked whether this potentiation effect of endogenously released CGRP extends more broadly across excitatory inputs in the BNST, by investigating the effect of high-frequency stimulation on spontaneous excitatory postsynaptic currents (sEPSCs). To do this, we first confirmed that BNST neurons received PBN_CGRP_ glutamatergic input by testing for oEPSCs in cells patched with a low-chloride internal solution and held at -70 mV. Under these conditions, excitatory currents are readily detected as inward sEPSCs while inhibitory currents are largely masked. After recording baseline spontaneous activity, we applied high-frequency optical stimulation (10 Hz, 5 s) to drive neuropeptide release from PBN_CGRP_ terminals in the BNST. We then measured sEPSCs again, in the 100 second window shown to potentiate oEPSCs, to assess whether CGRP release modulated spontaneous excitatory synaptic activity.

Example traces of spontaneous postsynaptic currents (sEPSCs) from male and female BNST neurons are shown in Figure 5A. There was no change in median sEPSC amplitude (Fig. 5B; male: p = 0.09, t = 1.84, n = 12 neurons; female: p = 0.22, t = 1.32, n = 9 neurons; paired t-test) or the distribution of sEPSC amplitudes in either sex (Fig. 5C; male: p > 0.99, D = 0.088; female: p = 0.18, D = 0.265, Kolmogorov–Smirnov test). There was also no change in median sEPSC frequency (Fig. 5D; males: p = 0.11, W = 42.00, n = 12 neurons; females: p = 0.20, W = –23.00, n = 9 neurons; Wilcoxon signed-rank test) or the distribution of interevent intervals following neuropeptide release (Fig. 5E; males: p = 0.74, D = 0.167; females: p = 0.65, D = 0.182; Kolmogorov-Smirnov test) in either sex. Taken together, these results indicate that CGRP release does not alter the amplitude or frequency of spontaneous excitatory postsynaptic currents in BNST neurons from either male or female mice, suggesting that while CGRP modulates PBN-specific glutamatergic inputs to BNST neurons, basal excitatory synaptic transmission is unaffected.

**Figure 5:**
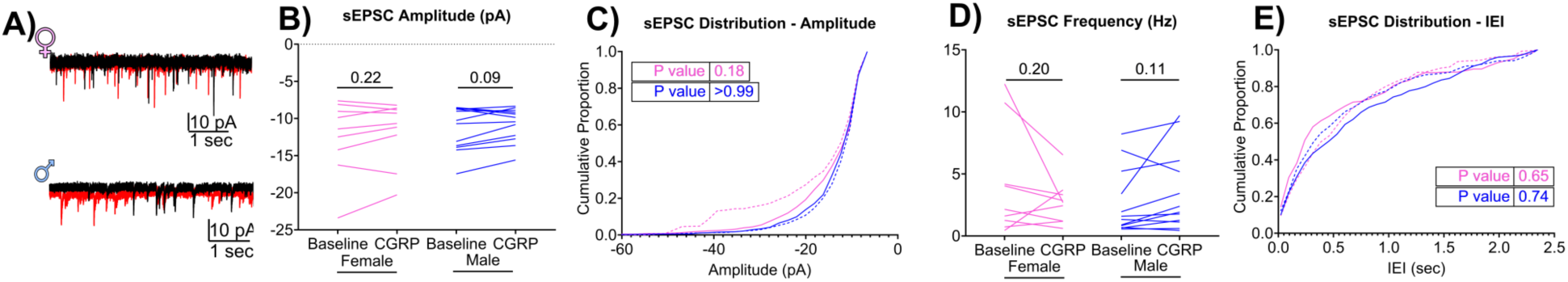
PBN CGRP does not alter spontaneous glutamate signaling. A) Example traces of spontaneous excitatory postsynaptic currents (sEPSCs) recorded from BNST neurons in female (♀) and male (♂) mice receiving PBN_CGRP_ glutamatergic input. Recordings used a high-chloride internal solution. Baseline (black) and post-high frequency stimulation (10 Hz, 5 s; red) activity are shown. B) Median sEPSC amplitude did not change during the first 100 s post-stimulation. C) Cumulative distributions of sEPSC amplitudes were unchanged from baseline (solid lines) following high-frequency stimulation (dashed lines) in both sexes. D) Median sEPSC frequency was unchanged in both sexes. E) Cumulative distributions of sEPSC interevent intervals (IEIs) were unchanged from baseline (solid lines) following high-frequency stimulation (dashed lines) in both sexes. Pink and blue lines or datapoints represent female or male neurons, respectively.

### No Sex Differences in PBN_CGRP_ Glutamatergic input to BNST Neurons

In addition to CGRP, PBN_CGRP_ inputs to the BNST also release glutamate. Sex differences in the strength and targeting of these glutamatergic inputs could contribute to sex differences in PBN activation of the BNST in pain and aversion. We investigated whether there are sex differences in the synaptic properties of PBN_CGRP_ glutamatergic inputs to the BNST. We first compared, in males and females, the proportion and distribution of these inputs.

oEPSCs were evoked in most neurons recorded in the terminal eYFP^+^ field in BNST (Fig. 6A) of both males (52 of 95 neurons) and females (40 of 59). There was no sex difference in the proportion of BNST neurons responsive to PBN_CGRP_ inputs (p = 0.11, χ² = 2.58; Chi-square test). Neurons demonstrating evoked oEPSCs were primarily located in the dorsolateral, juxtacapsular, and ventral oval nuclei of the BNST (Fig. 6B) in both sexes, consistent with the pattern of the PBN_CGRP_ terminal field (Fig. 3B). The minority of BNST neurons that did not receive PBN_CGRP_ glutamatergic input were recorded from within these same regions (Fig. 6B). These data suggested there is no sex difference in the proportion or subnuclear localization of BNST neurons receiving PBN_CGRP_ glutamatergic inputs.

**Figure 6:**
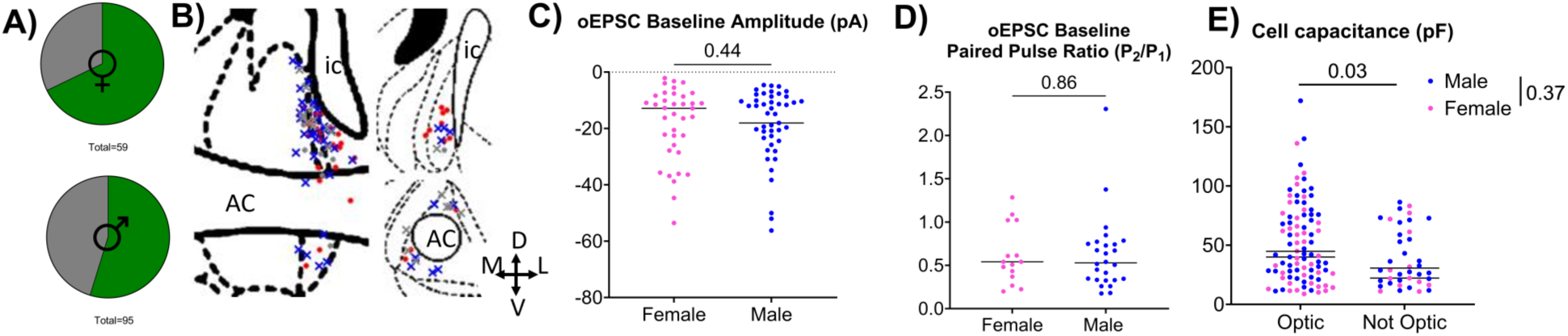
No sex differences in glutamate signaling from PBN_CGRP_ inputs to the BNST. A) The proportion of recorded BNST neurons exhibiting detectable optically evoked excitatory postsynaptic currents (oEPSCs) in response to optical stimulation of ChR2-expressing PBN_CGRP_ inputs was similar in males (♂) and females (♀) (p = 0.11, χ² = 2.58; Chi-square test). B) Locations of recorded BNST neurons in males (x’s) and females (dots). Colored symbols (blue = male, red = female) denote neurons with detectable oEPSCs; gray symbols indicate neurons without detectable oEPSCs). Coronal sections are shown at Bregma +0.38 mm (bottom right), +0.20 mm (left), and +0.02 mm (top right). White matter landmarks include the anterior commissure (AC) and internal capsule (ic). Compass indicates medial (M), lateral (L), dorsal (D), and ventral (V). C) oEPSC amplitudes and D) paired-pulse ratios (PPR, pulse 2 / pulse 1 amplitude, delivered with 150 msec inter-stimulus interval) were not different between male and female BNST neurons. E) BNST neurons that did not respond to optically-evoked PBN^CGRP^ glutamate release (optic non responders) inputs had lower cell capacitance compared to optic responders. Two-way ANOVA revealed there was no effect of sex or interaction effect.

To test for sex differences in PBN_CGRP_ glutamatergic signaling strength in the BNST, we compared oEPSC amplitude and paired pulse ratios between males and females. There was no sex difference in baseline oEPSC amplitudes (Fig. 6C, p = 0.44, U = 817; n = 49 male, 37 female neurons; Mann-Whitney test). This suggests that input strength of PBN_CGRP_→BNST glutamatergic synapses is similar in males and females. There was also no sex difference in baseline PPR (Fig. 6D, p = 0.86, U = 188; n = 26 male, 15 female neurons; Mann-Whitney test). This suggests that PBN_CGRP_ glutamatergic release probability is similar between males and females. Additionally, the oEPSC PPR was less than 1 in 35 / 41 BNST neurons patched (Fig. 6D). This indicates that PBN_CGRP_ glutamatergic inputs to the BNST exhibit paired pulse depression; a PPR below one is associated with a high probability of neurotransmitter release.

Taken together, there were no sex differences in the proportion or signaling strength of PBN_CGRP_ glutamatergic inputs to the BNST. However, the BNST is a heterogeneous structure, comprising of diverse neuron types with distinct electrophysiological (Hammack et al., 2007; Hazra et al., 2011; Miura et al., 2023; Rodríguez-Sierra et al., 2013) and molecular characteristics (Lebow and Chen, 2016; Ortiz-Juza et al., 2021). Many of these subpopulations are linked with divergent functions in aversion processing (Ortiz-Juza et al., 2021). Therefore, we tested whether there are sex differences in the BNST neurons that receive PBN_CGRP_ inputs, as sex differences in the targets of PBN_CGRP_ neurons could influence the downstream functional effects of CGRP-dependent glutamate potentiation.

To investigate this, we compared the intrinsic properties of female and male BNST neurons that receive PBN_CGRP_ inputs. There were no differences in access resistance (*p* = 0.99, U = 948; *n* = 50 male, 38 female neurons; Mann-Whitney test), baseline holding current (*p* = 0.33, U = 142; *n* = 22 male, 16 female neurons; Mann-Whitney test), membrane resistance (*p* = 0.47, U = 886.5; *n* = 50 male, 39 female neurons; Mann-Whitney test), or cell capacitance (*p* = 0.18, U = 811; *n* = 50 male, 39 female neurons; Mann-Whitney test) suggesting that passive membrane properties are similar between sexes. Because access resistance is influenced by the quality of the electrical seal with the patched cell, and cell capacitance influences signal-to-noise ratio, these findings increase our confidence that underlying sex differences in glutamatergic signaling strength were not overlooked due to technical limitations.

There were no differences in access resistance or membrane resistance between BNST neurons that responded to PBN^CGRP^ glutamate release (“optic responders”), identified by the presence of an oEPSC following optical stimulation of ChR2-expressing PBN^CGRP^ terminals, and those that did not (“optic non-responders”). Access resistance was not influenced by oEPSC response, sex, or the interaction (oEPSC response: *p* = 0.78, F(1,126) = 0.08; sex: *p* = 0.65, F(1,126) = 0.21; interaction: *p* = 0.58, F(1,126) = 0.31; two-way ANOVA) nor was membrane resistance (oEPSC response: *p* = 0.12, F(1,127) = 2.50; sex: *p* = 0.44, F(1,127) = 0.59; interaction: *p* = 0.78, F(1,127) = 0.08; two-way ANOVA).

However, there was an effect of response state on cell capacitance (Fig. 6E, oEPSC response: p = 0.03, F(1,127) = 5.06), with no effect of sex (p = 0.37, F(1,127) = 0.80) or interaction (p = 0.74, F(1,127) = 0.11; two-way ANOVA). Optic responders had a higher mean cell capacitance (52.35 pF ± 5.52(SD)) compared to optic non-responders (38.11 pF ± 2.50 (SD)). This suggests that BNST neurons receiving PBN_CGRP_ glutamatergic input, regardless of sex, tend to have higher cell capacitance than those that do not. Since cell capacitance is influenced by membrane surface area, this could indicate that input-recipient BNST neurons are larger than non-recipient cells. This difference may also reflect a technical factor, such as greater preservation of dendritic processes during slice preparation in responding neurons.

Together, these findings indicate that PBN_CGRP_ glutamatergic inputs to the BNST are similar in males and females in terms of their prevalence, strength, and synaptic properties. However, neurons that receive these inputs exhibit higher cell capacitance than non-responsive neurons, suggesting they may represent a distinct, possibly larger, subpopulation within the BNST.

### Heterogeneity in CGRP’s Effect on Evoked Responses

We further explored the heterogeneous effect of both endogenous and exogenous CGRP on evoked inhibitory (Fig. 3K) and excitatory (Fig. 4E) responses. Given that different subnuclei of the BNST are associated with distinct functions (Gungor and Pare, 2016), we investigated whether neurons within subregions of the anterolateral BNST respond differently to CGRP. We grouped neurons into three categories: Where CGRP increased (>120% of baseline), decreased (<80% of baseline), or caused no change in response amplitude. For experiments with endogenous (optically evoked) CGRP release, categories were based on the normalized oEPSC amplitude within 100 seconds of optic tetanus. For bath CGRP application, categories were based on the normalized IPSC amplitude 6-12 minutes after CGRP application. These categories were plotted onto a BNST atlas based on a 4x photo taken of the recorded neuron’s location. There was no spatial clustering in neurons responses to either endogenous (Fig. 7A) or exogenous (Fig. 7B) CGRP, suggesting that a neuron’s response profile to CGRP cannot be predicted by its location.

**Figure 7:**
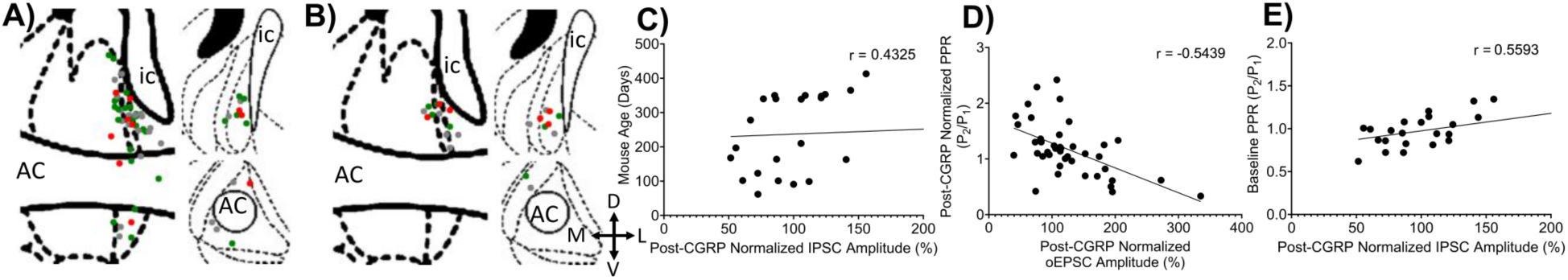
Heterogeneity in CGRP’s Effect on Evoked Responses in BNST neurons. A, B) Coronal maps showing locations of recorded BNST neurons that exhibited increased (green), decreased (red), or unchanged (gray) responses (defined as ±20% from baseline) in either A) optically evoked EPSC (oEPSC) amplitude following high-frequency stimulation, or B) evoked IPSC amplitude following exogenous CGRP (500 nM) application. Slices are shown at Bregma +0.38 mm (bottom right), +0.20 mm (left), and +0.02 mm (top right). White matter tracts include the anterior commissure (AC) and internal capsule (ic). Orientation: medial (M), lateral (L), dorsal (D), and ventral (V). C) A weak positive correlation was observed between the age of the mouse and normalized IPSC amplitude following CGRP application. D) A modest negative correlation was observed between normalized oEPSC potentiation amplitude and normalized paired-pulse ratio (PPR) following high-frequency optic stimulation. E) A modest positive correlation was observed between baseline IPSC PPR and normalized IPSC amplitude following CGRP application (500 nM).

BNST neurons can be classified based on electrophysiological parameters (Hammack et al., 2007; Hazra et al., 2011; Miura et al., 2023; Rodríguez-Sierra et al., 2013). To test whether certain CGRP response profiles are seen in functionally-distinct neurons, we conducted an exploratory correlation analysis. We compared either the normalized oEPSC amplitude within 100 seconds of optic tetanus, or the normalized IPSC amplitude 6 to 12 minutes after CGRP application, to various experimental factors and neuron properties. These comparisons were grouped into three categories: experimental parameters, passive membrane properties, and baseline input strength. The results of these correlations are summarized in Tables 2-4.

We first examined experimental parameters: animal age, viral incubation time, lateralization (right vs. left side), and access resistance. There was no correlation between the normalized post-stimulus oEPSC amplitude and any of these parameters (Table 1), except for a positive correlation between post-CGRP IPSC amplitude and mouse age (Table 1, Fig. 7C, r = 0.43, p = 0.039; n = 23 neurons; Spearman correlation). This suggests a weak correlation between age and the inhibitory response to CGRP, where older mice show a greater increase in inhibitory tone following CGRP release, consistent with reported age-related changes in neuronal excitability (Smithers et al., 2017).

We then explored whether passive membrane properties, including baseline holding current, membrane resistance, and cell capacitance, influenced oEPSC and IPSC responses to CGRP. There was no correlation between post-CGRP oEPSC amplitude or IPSC amplitude and any of these properties (Table 2). Since variations in membrane resistance and capacitance can affect the accuracy of the patch-clamp measurements, the lack of a correlation between these properties and the oEPSC or IPSC amplitudes after CGRP application increases our confidence that heterogenous responses were not simply the result of technical limitations between recorded neurons.

Finally, we tested whether baseline glutamatergic or GABAergic input strength correlated with post-CGRP oEPSC or IPSC responses. There was no correlation between CGRP response and baseline glutamatergic oEPSC amplitude, indicating that the strength of PBN_CGRP_ glutamatergic inputs does not predict post-CGRP oEPSC response. Similarly, baseline GABAergic input strength (IPSC amplitude) did not correlate with post-CGRP IPSC response (Table 3).

However, there was a correlation between CGRP responses and PPR. Normalized oEPSC potentiation amplitude was inversely correlated with normalized post-CGRP PPR (Fig. 7D, r = -0.54, p = 0.0002; n = 41 neurons; Spearman correlation). This suggests that neurons showing oEPSC potentiation after endogenous CGRP release tend to exhibit a concurrent reduction in PPR, indicating enhanced presynaptic glutamate release from PBN_CGRP_ terminals. While the overall PPR for the population was unaffected by glutamate release (Fig. 4I), these findings imply that a presynaptic mechanism may be engaged in those neurons exhibiting the greatest increases in potentiated oEPSC signaling following CGRP release. This is unlikely to result from autocrine CGRP signaling, as PBN CGRP neurons do not express the CGRP receptor (Oliver et al., 1998; Pauli et al., 2022; Van Rossum et al., 1997). Instead, presynaptic facilitation may arise from retrograde signaling, such as reduced tonic endocannabinoid activity (Lovinger, 2008) or increased nitric oxide signaling (Hardingham et al., 2013). Indeed, BNST nitric oxide is implicated in sympathetic hyperactivation (Alves et al., 2009; Barretto-de-Souza et al., 2018; Oliveira et al., 2018) and anxiogenic effects mediated through glutamatergic, NMDA-dependent pathways (Faria et al., 2016; Hott et al., 2017).

On the inhibitory side, post-CGRP IPSC amplitude was correlated with baseline inhibitory PPR (Fig. 7E, r = 0.56, p = 0.0055; n = 23; Spearman correlation). This suggests that neurons with weaker presynaptic release probability at baseline (evidenced by paired-pulse facilitation) are more likely to exhibit larger potentiation of IPSC amplitude after CGRP application. These results imply that CGRP may selectively potentiate weaker inhibitory inputs to BNST neurons.

Overall, there was no spatial clustering of CGRP response profiles within the anterolateral BNST, nor any correlations with experimental parameters, membrane properties, or baseline input strength. However, correlations with PPR implicate projection-specific contributions to the variability in CGRP responses.

## Discussion

Our findings demonstrate that PBN_CGRP_ neurons release neuropeptides in the BNST in response to aversive stimuli in a sex-independent manner. These PBN_CGRP_ neurons also provide glutamatergic inputs to the BNST, with no sex differences in input prevalence, strength, or synaptic properties. In both sexes, high-frequency activation of PBN_CGRP_ terminals induces endogenous CGRP release that transiently potentiates PBN_CGRP_→BNST glutamatergic signaling, without increasing overall spontaneous excitatory tone. Whereas CGRP selectively increases spontaneous inhibitory signaling in males, evoked inhibitory signaling and presynaptic GABA release probability are not consistently modulated in either sex. Across measurements of evoked excitatory and inhibitory currents, CGRP effects are heterogeneous, with neurons exhibiting potentiation, suppression, or no change. This variability is not related to the location of the recorded neurons, but is correlated with input-specific presynaptic properties, such as paired-pulse ratio.

PBN CGRP neurons release neuropeptides in BNST during aversive processing In both males and females, applying an aversive stimulus—either foot shock or mechanical pinch in lightly anesthetized animals—increased neuropeptide release similarly in magnitude and duration. Response amplitude scaled with stimulus intensity, but no sex differences were observed.

Following neuropeptide release, both sexes exhibited a prolonged suppression of CybSEP2 fluorescence below baseline that persisted for at least 1 minute after stimulation. This signal suppression cannot be explained by the kinetics of the CybSEP2 sensor (Kim et al., 2024). We interpret this suppression as depletion of the readily recruitable pool of large dense-core vesicles (LDCVs) within PBN CGRP. Although CybSEP2 is a pH-sensitive sensor that fluoresces more strongly at alkaline pH, it retains weak fluorescence within the acidic intravesicular environment (pH 5–5.5, (Johnson and Scarpa, 1976)) of LDCVs prior to release. Therefore, a reduction in the readily releasable LDCV pool at PBN CGRP terminals may manifest as a decrease in baseline fluorescence. A transient reduction in available LDCVs following a release event is further supported by prior characterization of LDCV recycling dynamics using the CybSEP2 sensor (Kim et al., 2024). In that study, consecutive tail pinches evoked neuropeptide release from PBN CGRP terminals in CeLC only when delivered at 2–3 minute intervals, whereas a second tail pinch delivered within 1 minute failed to elicit a response, suggesting that LDCVs require more than 1 minute to be recycled and replenished at the terminal.

To our knowledge, ours is the first study examining PBN CGRP neuropeptide release in the BNST. Our findings are consistent with aversive stimuli-evoked release from PBN CGRP neurons in the central amygdala (Kim et al., 2024). Although nociceptive responses persist under anesthesia—as evidenced by autonomic changes, such as increased heart rate and blood pressure, in response to noxious stimuli (Cividjian et al., 2017) —the magnitude of PBN CGRP release we observed is likely reduced compared to awake conditions. This is consistent with our evidence that anesthesia suppresses, but does not eliminate, PBN neuronal activity in response to aversive input (Smith et al., 2023).

Real-time measurement of released neuropeptides is technically challenging due to their large size and low extracellular concentrations, but our findings are consistent with previous reports, showing that CGRP can be released following neuronal activation (Wang et al., 2019) or in response to painful stimuli (Frese et al., 2021). These studies relied on a single measurement or on peripheral blood sampling. By using CybSEP2 as a proxy for LDCV release, we quantified, in real time and with high spatial resolution, the dynamics of evoked neuropeptide release, allowing direct comparisons between males and females.

A limitation of our approach is that it cannot distinguish among co-expressed neuropeptides. PBN CGRP neurons co-express several neuropeptides, including pituitary adenylate cyclase-activating polypeptide (PACAP), substance P, neurotensin, and corticotropin-releasing hormone (Pauli et al., 2022; Sanz et al., 2009), many of which have been linked to aversive processing (Boucher et al., 2021; Hsieh et al., 2020; Jiang et al., 2023; Kainu et al., 1993; Seiglie et al., 2023; Kelly S. Sink et al., 2013; K.S. Sink et al., 2013; Sink et al., 2011). To our knowledge, no studies have examined the release kinetics of these neuropeptides during aversive processing.

Together, these findings provide the first direct, real-time evidence that PBN CGRP neurons release neuropeptides in the BNST during aversive processing in a stimulus-dependent, but sex-independent manner.

### CGRP biases female BNST toward hyperexcitability

Both CGRP signaling and the BNST are causally implicated in aversive behaviors, including chronic pain (Huang et al., 2021; Maruyama et al., 2018; Minami, 2019; Minami and Ide, 2015; Kelly S. Sink et al., 2013; Sink et al., 2011; Song et al., 2020; Uchida et al., 2024; Yamauchi et al., 2022). Moreover, activation of BNST neurons that receive parabrachial inputs modulates fear- and anxiety-like behaviors in a sex-dependent manner in stress-relevant contexts (Jaramillo et al., 2020; Van Doorn et al., 2025), yet, whether CGRP signaling within this pathway contributes to these differences remains unknown. Because our findings and others’ support sex-dependent effects of CGRP in other brain regions (Lorsung et al., 2025; Paige et al., 2022) we tested the alternative hypothesis that CGRP exerts sex-dependent effects on its downstream targets in BNST.

### Inhibitory signaling

CGRP did not affect inhibitory synaptic strength: neither optically evoked endogenous PBN CGRP release nor exogenous CGRP altered GABA-mediated currents evoked by electrical stimulation. This contrasts with prior work demonstrating CGRP-mediated potentiation of postsynaptic GABA responses in rat BNST (Gungor and Pare, 2014). Differences in species and developmental stage (≥8-week-old mice here vs. 4 to 7-week-old rats in Gungor and Pare, 2014) may contribute to this discrepancy, as CGRP receptor expression declines with age in other regions, such as the spinal cord (Qin et al., 2024).

In contrast to its lack of effects on evoked GABAergic currents, CGRP increased the frequency of spontaneous GABAergic currents in males, but not females, without altering their amplitude in either sex. These findings suggest that CGRP increases GABA release in males without altering postsynaptic responsiveness to GABA. Together with the lack of effect on evoked PPR, this is consistent with a mechanism in which CGRP increases the excitability of a local inhibitory neuronal population, as has been observed in the central amygdala (Han et al., 2010). This mechanism aligns with reports of pain-induced increases in excitability of inhibitory BNST subpopulations, largely characterized in male mice or in studies where sex was not specified (Huang et al., 2021; Uchida et al., 2024; Yamauchi et al., 2022). Also relevant are findings that aversive processing, such as morphine withdrawal (Luster et al., 2020) or fear conditioning (Hon et al., 2025), are associate with increased GABA signaling in males, but decreased signaling in females. Together, our results identify CGRP as a rapid, sex-specific modulator of intra-BNST inhibitory tone (Nagano et al., 2015), preferentially enhancing presynaptic GABA release in males.

### Excitatory signaling

Endogenous CGRP release transiently potentiated PBN CGRP glutamatergic inputs to BNST neurons in both sexes. These results contrast with our findings in the central amygdala, where endogenously released CGRP induced potentiation of PBN CGRP glutamatergic inputs preferentially in females (Lorsung et al., 2025). In the BNST, potentiation of glutamatergic signaling was observed at PBN_CGRP_ inputs, while spontaneous excitatory postsynaptic currents were unchanged. This indicates that CGRP modulation does not manifest as a global increase in excitatory tone.

Thus, CGRP potentiates excitatory PBN inputs in both sexes, but selectively enhances inhibitory tone in males. This male-specific enhancement of inhibitory tone may act as a compensatory mechanism to limit BNST excitability, potentially contributing to sex differences BNST hyperexcitability under conditions of elevated CGRP release, including pain, stress, and anxiety.

## Acknowledgments

This work was supported by National Institutes of Health–National Institute of Neurological Disorders and Stroke Grants R01NS099245, R01NS069568, R01NS127827 and Fellowship F31NS134126 (to RL). The content is solely the responsibility of the authors and does not necessarily represent the official views of the National Institutes of Health. The authors thank the University of Maryland School of Medicine Virus Vector Core (Baltimore, MD) for viral vector production.

## Notes

### Competing Interest Statement

The authors have declared no competing interest.

